# Cocaine place conditioning strengthens location-specific hippocampal inputs to the nucleus accumbens

**DOI:** 10.1101/105890

**Authors:** Lucas Sjulson, Adrien Peyrache, Andrea Cumpelik, Daniela Cassataro, György Buzsáki

## Abstract

Conditioned place preference (CPP) is a widely used model of addiction-related behavior whose underlying mechanism is not understood. In this study, we used dual site silicon probe recordings in freely moving mice to examine interactions between the hippocampus and nucleus accumbens in cocaine CPP. We found that CPP was associated with recruitment of nucleus accumbens medium spiny neurons to fire in the cocaine-paired location, and this recruitment was driven predominantly by selective strengthening of hippocampal inputs arising from place cells that encode the cocaine-paired location. These findings provide in vivo evidence that the synaptic potentiation in the accumbens caused by repeated cocaine administration preferentially affects inputs that were active at the time of drug exposure. This provides a potential physiological mechanism by which drug use becomes associated with specific environmental contexts.

## INTRODUCTION

Drug addiction is a debilitating condition for which few effective treatments exist, in large part because the underlying mechanisms are not well understood. A key insight into the nature of addiction is that drug use becomes associated with the environmental context in which the drug was administered. Subsequent re-exposure to the associated context then leads to cravings or drug-seeking behavior. One of the simplest animal models of this association is cocaine conditioned place preference (CPP), in which cocaine is repeatedly paired with a specific spatial location causing the animal to spend more time in that location during subsequent exploration. However, despite its simplicity and relevance, the underlying mechanistic basis of cocaine CPP is still not understood.

The nucleus accumbens (NAc) is a part of the ventral striatum believed to play a central role in reward- and addiction-related behaviors including CPP. Medium spiny neurons (MSNs) in the NAc are known to fire preferentially near sites where overtrained animals collect water or food rewards (Lansink et al., 2008; Lavoie and Mizumori, 1994; Miyazaki et al., 1998; van der Meer et al., 2010), providing a potential substrate for location-reward association. However, German et al. (German and Fields, 2007) found that morphine place conditioning led to *less* MSN firing in the morphine-paired location, suggesting that the association of drug reward and spatial location may occur through different mechanisms. Nevertheless, focal lesions (Kelsey et al., 1989) or D1 antagonist injections (Baker et al., 1998) in the NAc block CPP acquisition, and focal amphetamine injections into the NAc are sufficient to induce CPP (Carr and White, 1983), suggesting that location-dependent MSN activity may drive CPP behavior.

A large glutamatergic input to the NAc arises from the CA1 and subiculum regions of the hippocampus (HPC) (Phillipson and Griffiths, 1985), which contain “place cells" that fire selectively in specific spatial locations (Kim et al., 2012; O’Keefe, 1976) or contexts (Komorowski et al., 2013). Simultaneous HPC-NAc recordings in rats suggest that hippocampal inputs carry spatial information to the NAc (Lansink et al., 2008; Lansink et al., 2009; Tabuchi et al., 2000; van der Meer and Redish, 2011a), and anatomical disconnection experiments indicate that hippocampus-accumbens interactions are necessary for CPP (Ito et al., 2008). However, the mechanism by which these inputs could mediate reward location-dependent MSN activity and CPP behavior is not known.

One possibility is that synaptic plasticity of hippocampal inputs to the NAc stores information about which locations were previously paired with cocaine. Systemic NMDA receptor blockade prevents CPP acquisition (Cervo and Samanin, 1995), suggesting that synaptic plasticity is necessary for CPP, and *ex vivo* slice experiments have shown that repeated cocaine exposure potentiates hippocampal synapses onto D1-positive MSNs in the NAc (Britt et al., 2012; Pascoli et al., 2014). Plasticity at corticostriatal synapses generally requires presynaptic activity (Calabresi et al., 1999), raising the possibility that cocaine may selectively strengthen the synapses that were most active at the time of drug exposure. Since CPP entails cocaine exposure in a specific location, we hypothesized that cocaine would preferentially strengthen NAc inputs from hippocampal place cells that encode the cocaine-paired location. To test this hypothesis, we performed simultaneous dual site silicon probe recordings in the HPC and NAc of freely moving mice in a cocaine CPP paradigm. Our findings indicate that cocaine conditioning recruits location-specific MSN firing, and this activity is driven predominantly by preferential strengthening of hippocampal inputs that encode the cocaine-paired location.

## RESULTS

### Dual site silicon probe recording during cocaine conditioned place preference

To measure functional interactions between the hippocampus and NAc in cocaine CPP, we performed simultaneous silicon probe recordings in both structures during CPP behavior (*n* = 6 mice) (Fig. 1A). We used established waveform and bursting criteria to identify putative pyramidal cells (PYRs) (Bartho et al., 2004; McCormick et al., 1985) in the hippocampus (*n* = 561 PYRs) and medium spiny neurons (MSNs) and interneurons (INs) in the NAc (*n* = 1293 MSNs, 313 INs; **Fig. S1)** (Schmitzer-Torbert and Redish, 2008; Yamin et al., 2013). Recordings were performed before (PRE) and after (POST) animals underwent cocaine place conditioning in a rectangular arena with a removable barrier (Fig. 1B, **Fig. S2**). We recorded 1-3 daily 30-minute PRE sessions prior to five consecutive days of cocaine conditioning, then 1-5 daily 30-minute POST sessions. This conditioning paradigm induced robust CPP in POST sessions (PRE = 27% in cocaine zone, POST = 51%, Wilcoxon rank sum test, *n* = 12 PRE, 15 POST sessions, *P* = 1.6 × 10^−6^; Fig. 1C, **Table S1**), which was attributable to an increase in the proportion of time spent immobile in the cocaine zone (**Fig. S3**).

**Figure 1:**
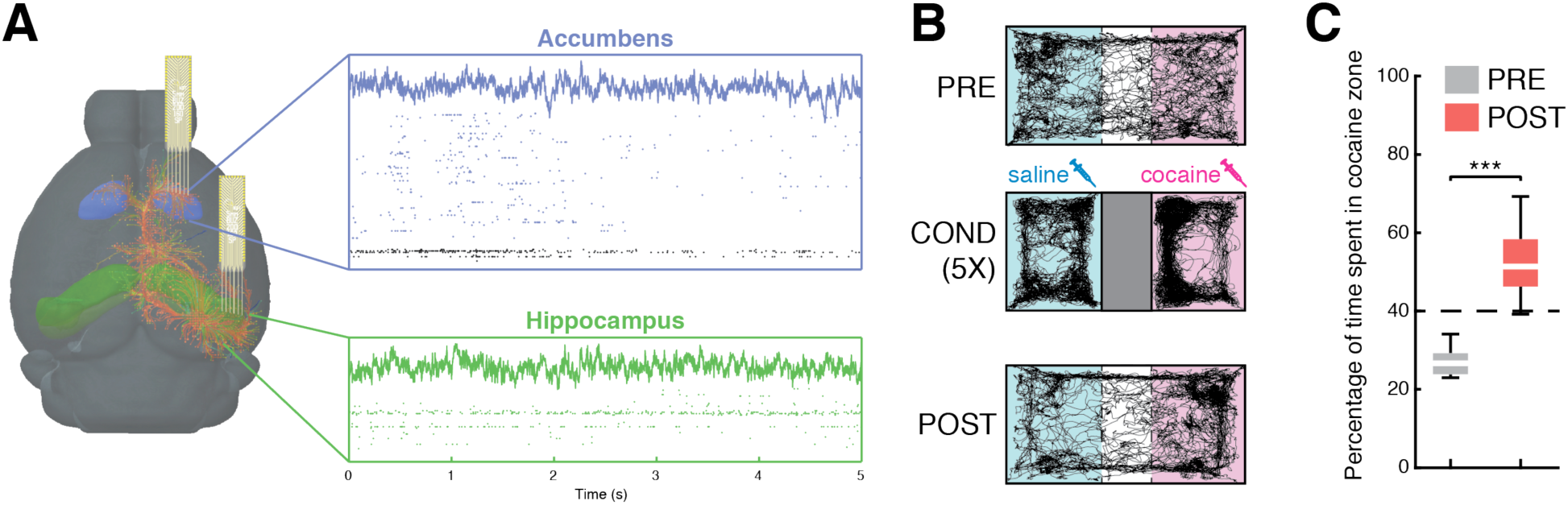
Cocaine conditioned place preference with dual-site silicon probe recording. **A)** Silicon probes were implanted in the hippocampus (green) and accumbens (blue), yielding LFP and single unit recordings (*n* = 1606 accumbens neurons, 725 hippocampal neurons). B) Mice (*n* = 6) were conditioned for five days with saline and cocaine 15 mg/kg IP. **C)** In POST sessions, animals exhibited a preference for the cocaine-paired zone (\*\*\**P* < 10^−5^, Wilcoxon sign rank test).

### Cocaine conditioning selectively increases MSN firing in the cocaine zone

We found that after cocaine conditioning, NAc MSNs fire at higher rates when the animal is in the cocaine zone than the saline zone (Cocaine index = -0.009 ± 0.011 PRE, 0.094 ± 0.013 POST, *n* = 545, 748 MSNs, unpaired t-test, *P* = 4.6 × 10^−9^; Fig. 2A). This effect was not seen in NAc INs (*n* = 162, 151 INs, unpaired t-test, *P* = 0.88, Fig. 2B), and for hippocampal PYRs a trend was noted that failed to reach statistical significance (*n* = 184, 377 PYRs, *P* = 0.11, Fig. 2C). However, the density of PYR place fields was higher in the cocaine zone in POST sessions (*P* = 2.6 × 10^−8^, **Fig. S6A**). Further, we found that the strength of behavioral CPP expression in a given POST session is correlated with the extent of increased MSN firing in the cocaine zone during that session (*n* = 15 POST sessions, R = 0.62, *P* = 0.01, Fig. 2D). This correlation occurs only in POST sessions and was not observed in NAc INs (R = 0.3, *P* = 0.27; Fig. 2E) or hippocampal PYRs (R = 0.1, *P* = 0.72; Fig. 2F). These observations support our hypothesis that cocaine CPP involves similar reward location-dependent MSN activity to that seen with non-drug rewards and prompted us to explore whether hippocampal inputs provide the location information utilized by MSNs.

**Figure 2:**
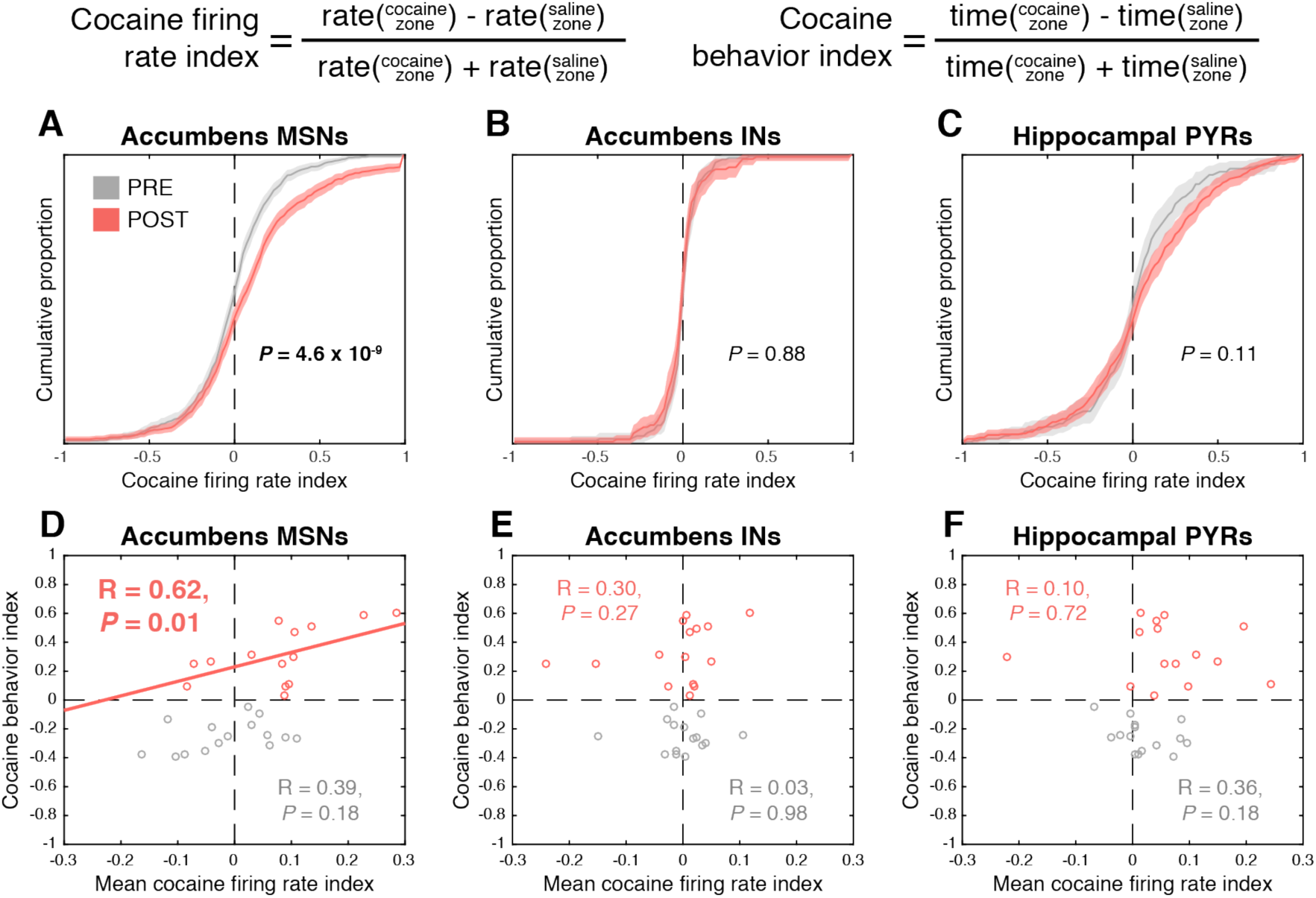
Cocaine conditioning selectively increases MSN activity in the cocaine zone. **A)** Accumbens MSNs fire preferentially in the cocaine zone after cocaine conditioning (*P* = 4.6 × 10^−9^). Accumbens putative interneurons (INs, **B**) and hippocampal pyramidal cells (PYRs, **C**) exhibit no shift after cocaine conditioning. **D)** Strength of behavioral CPP expression in each recording session is correlated with firing rate index in MSNs in POST (*P* = 0.01) but not PRE (*P* = 0.18) conditions. **E)** Behavioral CPP expression is not correlated with firing rate index in INs (*P* = 0.88 PRE, 0.27 POST). **F)** Behavioral CPP expression is not correlated with firing rate index in hippocampal PYRs (P = 0.18 PRE, 0.72 POST).

### MSNs exhibit signatures of decoding spatial location from hippocampal inputs

Although our CPP protocol was designed to ensure that CPP was hippocampus-dependent (Ito et al., 2008; Meyers et al., 2003) (**Fig. S2**), we first focused on analysis of spatial firing patterns of MSNs to verify that location-dependent MSN firing was driven by hippocampal inputs. Our initial analysis found that MSNs exhibit spatially biased firing patterns and also exhibit modulation by running speed (not shown). This complicates the analysis of MSN firing because CPP behavior fundamentally entails a strong correlation between movement and spatial location (**Fig. S3**). We attempted to disentangle this issue using generalized linear models (GLMs) and model cross-validation, finding that the best model fit included separate terms for location and speed modulation (**Table S2**). The GLM method enabled us to analyze these two phenomena separately and determine that MSNs exhibit running speed modulation that is independent of location and stronger than HPC PYRs (*n* = 1293 MSNs, 561 PYRs, Wilcoxon rank sum test, *P* = 3.9 × 10^−18^), which in turn are more strongly modulated than NAc INs (**Fig. S4a**, *n* = 561 PYRs, 313 INs, Wilcoxon rank sum test, *P* = 3.0 × 10^−5^). After correcting for running speed, MSNs encoded approximately as much spatial information (Skaggs et al., 1993) as hippocampal PYRs (Fig. 3B, *n* = 1293 MSNs, 561 PYRs, Wilcoxon rank sum test, *P* = 0.20) and significantly more than INs (*n* = 1293 MSNs, 313 INs, *P* = 1.1 × 10^−38^). We thus hypothesized that MSNs received spatial information via inputs from the hippocampus. To test this, we used GLMs to predict MSN spike trains from combinations of hippocampal activity and location (Fig. 3C). Using cross-validation, we found that adding hippocampal activity to a model containing no information other than baseline firing rate led to improvements in prediction quality. However, the improvement in prediction quality was smaller if we added hippocampal activity to a model that already contained information about location (Fig. 3D, *n* = 1203 MSNs, Wilcoxon signed rank test, *P* = 4.1 × 10^−11^). This is fully consistent with NAc MSNs decoding information about spatial location from hippocampal inputs. The prediction quality was highest when HPC led NAc by a time lag of ∼30 ms (Fig. 3E), suggesting that this effect represented information transfer from HPC to NAc rather than both structures receiving this information from a common input. A similar analysis on running speed found that MSNs decode information about running speed from hippocampal inputs as well (**Fig. S4**).

**Figure 3:**
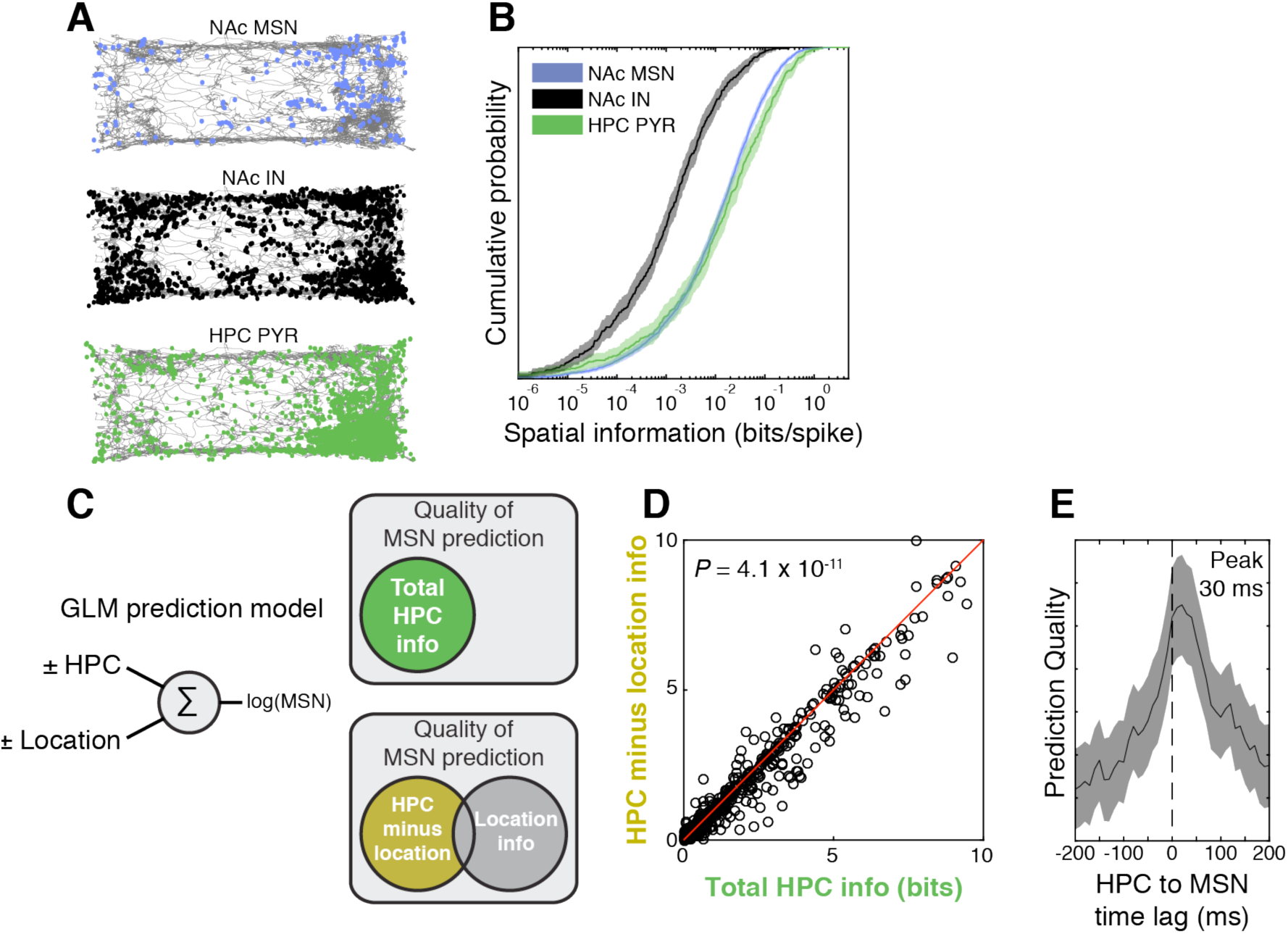
Accumbens MSNs decode spatial location from hippocampal inputs. **A)** Example session illustrating spiking activity (dots) of accumbens MSNs, INs and hippocampal pyramidal cells (HPC PYRs) in the testing apparatus. The gray line indicates the movement trajectory of the mouse. **B)** After correction for running speed, MSNs and PYR activity carries similar amounts of spatial information (*P* = 0.20 for MSN vs. PYR), but MSN activity carries more spatial information than IN activity (*P* = 1.0 × 10^−38^). **C)** GLMs enable prediction of individual MSN spike trains from the combination of PYR activity and location as predictors. **D)** Adding hippocampal activity as a predictor to a model already containing explicit location information results in a smaller improvement in prediction quality (*P* = 4.1 × 10^−11^) than for a model without explicit location information (note that most points are right of the diagonal). This suggests that predicting MSN spike trains from PYR inputs implicitly decodes information about spatial location. **E)** MSN spike train prediction quality is highest when PYR activity leads MSN activity by ∼30 ms, supporting the hypothesis that information is transferred from PYRs to MSNs.

### Assembly prediction analysis reveals increased hippocampus-accumbens coupling after cocaine conditioning

We next addressed the question of whether hippocampal inputs to the accumbens are strengthened by cocaine conditioning. To test this, we performed an assembly prediction analysis using PYR assemblies as predictors in a linear model that predicts the activity of a single MSN assembly (Fig. 4A, **Methods**). Consistent with our hypothesis that MSNs decode spatial information from hippocampal inputs, we found that the quality of the assembly prediction was correlated with the rate of spatial information encoded by the MSN assembly in POST, but not PRE, sessions (Fig. 4B). The spatial information rate was also higher in POST sessions (Fig. 4C, *n* = 34, 95 assemblies, Wilcoxon rank sum test, *P* < 0.05), suggesting that cocaine conditioning strengthens hippocampus-accumbens coupling.

**Figure 4:**
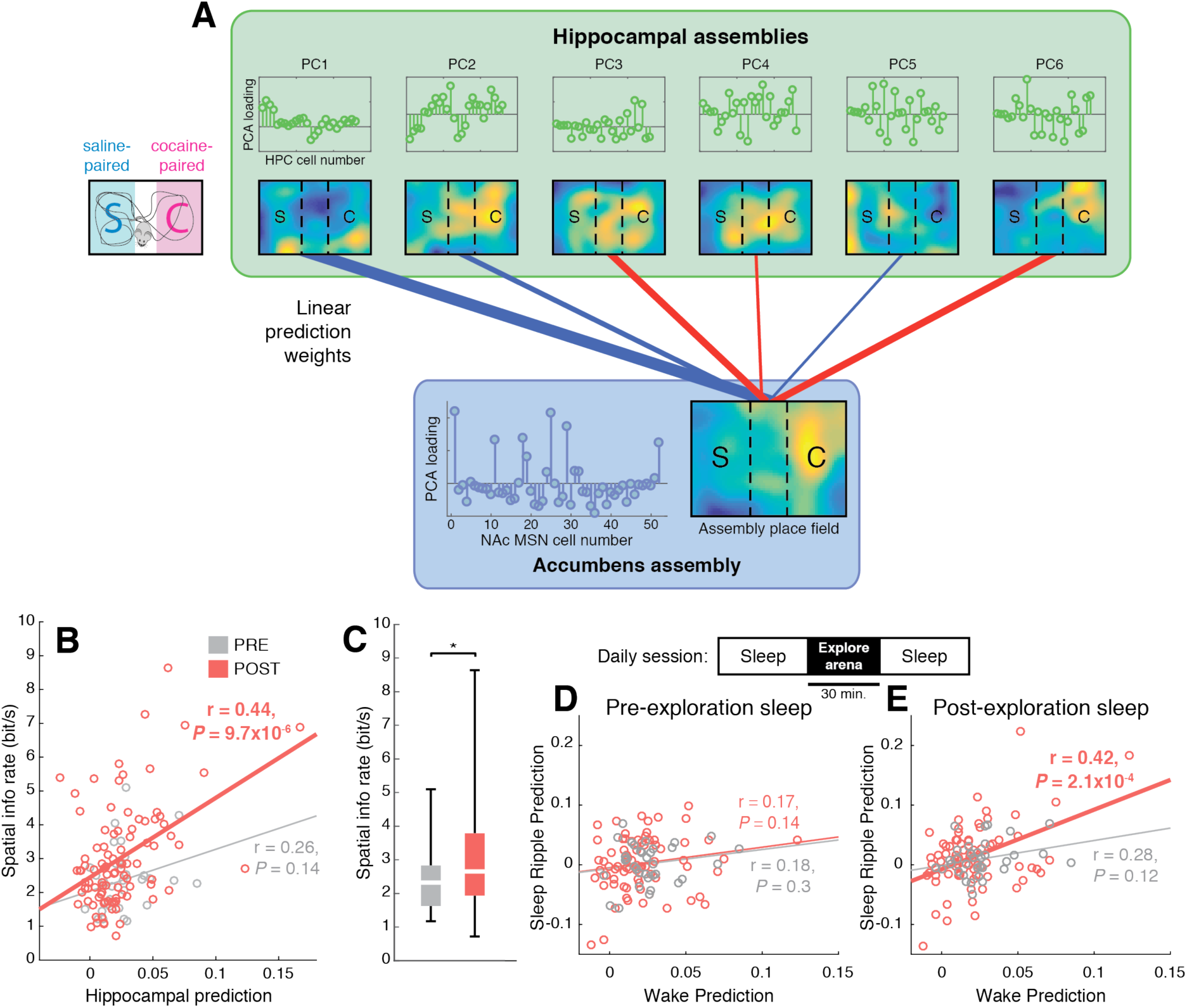
Cocaine conditioning increases the strength of functional coupling between hippocampal PYRs and accumbens MSNs. **A)** Activity of PYR assemblies predicts activity of MSN assemblies. **B)** The spatial information rate of an MSN assembly is correlated with the accuracy by which its activity can be predicted from PYR assemblies, in POST (R = 0.44, *P* = 9.7 × 10^−6^), but not PRE (R = 0.26, *P* = 0.14) sessions. **C)** Spatial information rate of MSN assemblies increases after cocaine conditioning (\**P* = 0.02). **D)** Model predictions for MSN assemblies during awake locomotion are uncorrelated with model predictions during sharp wave ripples in pre-exploration sleep. **E)** Model predictions during awake locomotion are correlated with predictions during sharp wave ripples in post-exploration sleep in POST (*P* = 2.1 × 10^−4^), but not PRE (*P* = 0.12), sessions. This suggests that cocaine conditioning strengthens coordinated hippocampus-accumbens replay.

To strengthen our finding that our assembly predictions were not due to a common input or sensory cues driving the hippocampus and accumbens, we performed an analysis of sleep replay events. Replay events, which occur during hippocampal sharp-wave ripple oscillations, consist of temporally compressed firing of place cells sequences that encode recently visited spatial locations (Lee and Wilson, 2002; Nadasdy et al., 1999; Skaggs and McNaughton, 1996) and are generated locally in the hippocampus (Buzsaki et al., 1983). We used data from awake exploration of the CPP arena to fit the prediction weights of the GLM, then tested model predictions both on awake data withheld from the training set and on data collected during sleep ripples occurring before and after the animal explored the CPP arena. We found that during pre-exploration sleep, when replay events encoding locations in the CPP arena do not occur, the sleep ripple prediction quality was uncorrelated with wake prediction quality (Fig. 4D). In contrast, during post-exploration sleep the sleep ripple prediction quality was significantly correlated with wake prediction quality in POST sessions (Fig. 4E), indicating strengthened hippocampus-NAc coupling.

### Hippocampally-modulated MSNs are preferentially recruited to fire in the cocaine zone

We next turned to the question of whether strengthened hippocampal inputs mediate the recruitment of additional MSN activity in the cocaine zone. To this end, we used simultaneous LFP recordings in the hippocampus to determine how strongly each MSN is phase-locked to the hippocampal theta oscillation. Several prior studies indicate that phase-locking with hippocampal theta is a marker for MSNs receiving strong hippocampal inputs (Jones and Wilson, 2005; Lansink et al., 2009; Tabuchi et al., 2000; van der Meer and Redish, 2011a). We found that MSNs with high theta modulation encode more spatial information than MSNs with low theta modulation (**Fig. S5A**) and are overrepresented among cells with either high and low cocaine indices (Fig. 5A, **Fig. S5B**). Dividing the MSNs into high-theta and low-theta halves, we found that the high-theta half shifts after conditioning toward encoding the cocaine zone (PRE 0.0082 ± 0.013, POST 0.087 ± 0.017, *n* = 261, 351 MSNs; Unpaired t-test, *P* = 2.7 × 10^−4^, Fig. 5B), while the low-theta half does not (PRE 0.0027 ± 0.016, POST 0.014 ± 0.016, *n* = 261, 351 MSNs; Unpaired t-test, *P* = 0.43, Fig. 5C). This suggests that strengthening of hippocampal inputs underlies recruitment of MSN firing to the cocaine zone.

**Figure 5:**
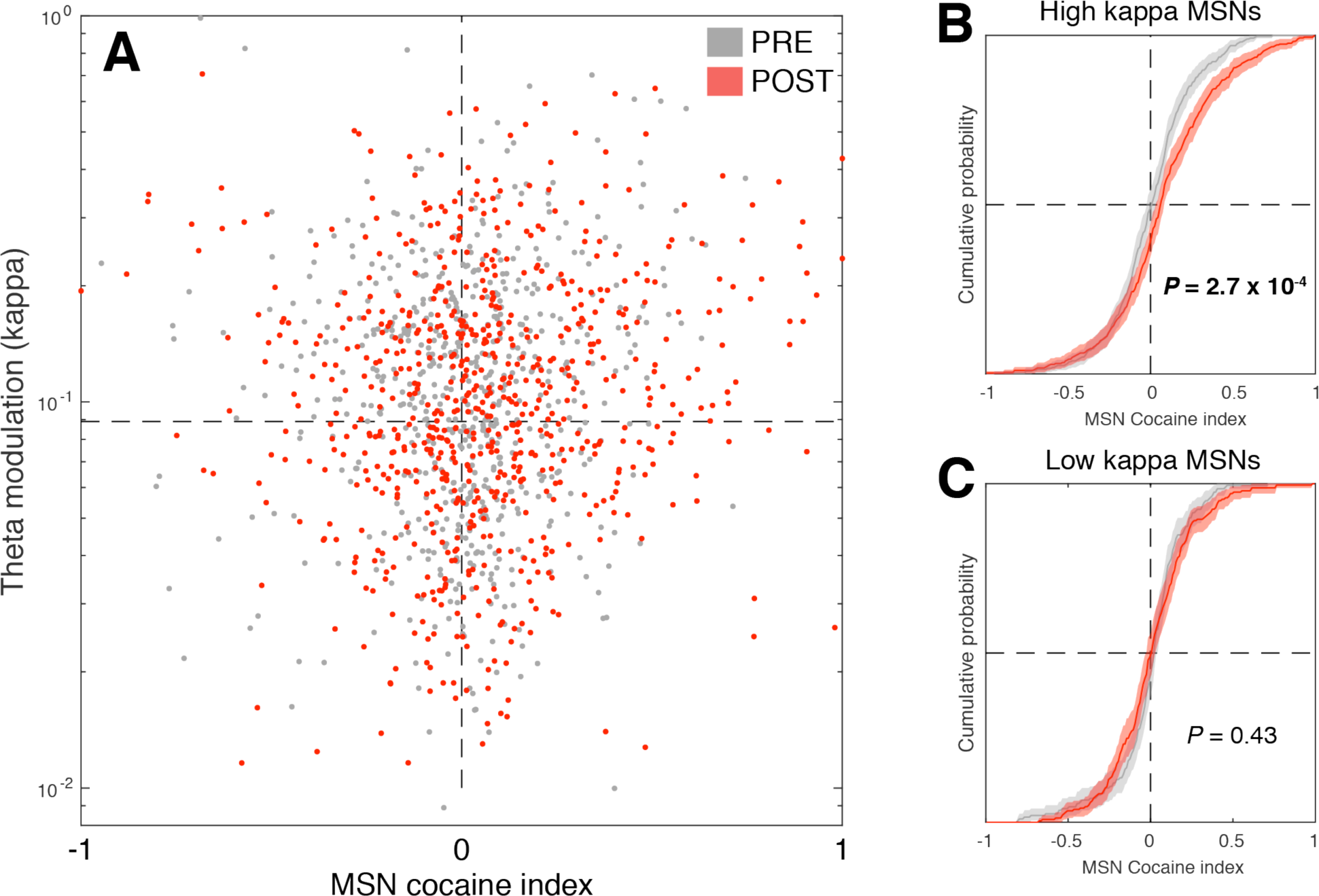
MSNs phase-locked to the hippocampal theta (∼8 Hz) oscillation show greater recruitment by cocaine conditioning. **A)** MSN cocaine firing rate index varies as a function of phase-locking to hippocampal theta. Note highest density of red dots (MSN POST) in the upper right quadrant. **B)** MSNs that are strongly phase-locked to hippocampal theta show significant increases in activity in the cocaine zone after conditioning (*P* = 2.7 × 10^−4^). **C)** MSNs that are weakly phase-locked to hippocampal theta do not show significant changes in cocaine zone activity after conditioning (*P* = 0.43).

### Selective strengthening of hippocampal inputs underlies MSN recruitment to the cocaine zone

To distinguish whether strengthening of hippocampal inputs is selective or nonselective (Fig. 6A), we fit a GLM to find connection weights that optimally predict MSN activity based solely on hippocampal activity (Fig. 6B). We then used this model with cross-validation to generate predictions of each MSN’s spike train, from which we calculated a cocaine index. Although the model contains no explicit location information, the cocaine indices predicted from hippocampal activity alone were significantly correlated with the observed cocaine indices (*n* = 531, 713 MSNs; PRE R = 0.45, *P* = 1.2 × 10^−7^; POST R = 0.59, *P* = 6.8 × 10^−68^; Fig. 6C), indicating that hippocampal activity is sufficient to predict MSN cocaine indices.

**Figure 6:**
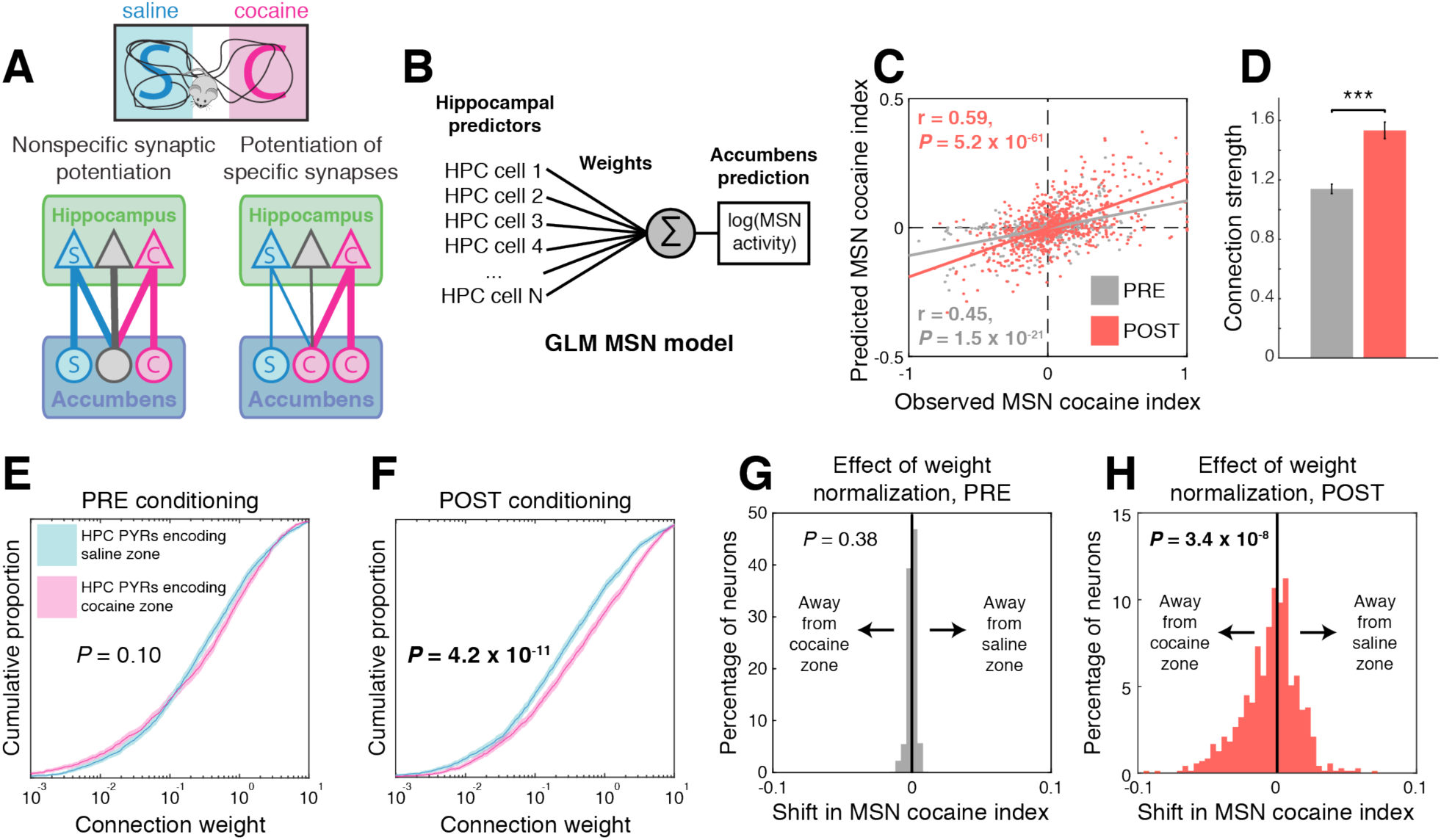
Cocaine place conditioning preferentially strengthens inputs from hippocampal pyramidal cells encoding the cocaine-paired location. **A)** Cocaine could either strengthen all hippocampal inputs equally or preferentially strengthen a subset of synapses. **B)** A GLM that predicts MSN spiking activity from PYR spiking activity can estimate connection weights and the effects of modifying them. **C)** Model-predicted MSN cocaine indices are significantly correlated with observed MSN cocaine indices (R = 0.45, *P* = 1.2 × 10^−7^ PRE; R = 0.59, *P* = 6.8 × 10^−68^ POST). **D)** Estimated connection weights are increased after cocaine conditioning (mean ± SEM, *P* = 1.2 × 10^−9^). **E)** In PRE sessions, connection weights from cocaine zone-encoding PYRs and saline zone-encoding PYRs have equal distributions (*P* = 0.10). **F)** In POST sessions, connections from cocaine zone-encoding PYRs have stronger weights than connections from saline zone-encoding PYRs (*P* = 4.2 × 10^−11^). **G)** In PRE sessions, adjusting connection weights from cocaine zone-encoding PYRs to match the distribution of weights from saline zone-encoding does not change predicted MSN cocaine indices (*P* = 0.38). **H)** In POST sessions, adjusting connection weights from cocaine zone-encoding PYRs significantly shifts predicted MSN cocaine indices away from the cocaine zone (*P* = 3.4 × 10^−8^), suggesting that the asymmetric connection weight distribution underlies increased MSN activity in the cocaine zone.

We next examined the PYR to MSN connection weights and found that, consistent with our earlier results, they increased in magnitude after cocaine conditioning (*n* = 4047, 3979 weights, two-tailed t test, *P* = 1.2 × 10^−9^; Fig. 6D). We then examined the strengths of connections from PYRs in the top and bottom quartile of cocaine indices, which encode the cocaine and saline zones, respectively. In PRE sessions there was no difference (*n* = 2011, 2036 weights, Wilcoxon rank sum test, *P* = 0.10; Fig. 6E), but in POST sessions connection weights arising from cocaine zone PYRs were significantly larger (*n* = 2014, 1981 weights, Wilcoxon rank sum test, *P* = 1.4 × 10^−6^; Fig. 6F), indicating that hippocampal inputs are strengthened selectively. To test whether the resulting asymmetry in the distribution of connection weights was responsible for the recruitment of MSN activity to the cocaine zone, we adjusted connection weights so that the distributions would be equal for PYRs encoding the cocaine and saline zones. In PRE sessions, this adjustment caused no change in MSN cocaine index (*n* = 531 MSNs, Wilcoxon signed rank test, *P* = 0.39, Fig. 6G). However, in POST sessions the MSN indices showed a significant shift away from the cocaine zone (*n* = 713 MSNs, Wilcoxon signed rank test, *P* = 3.4 × 10^−8^, Fig. 6H), indicating that the connection weight asymmetry contributes to MSN recruitment to the cocaine zone.

Since cocaine conditioning also increases the density of PYR place fields in the cocaine zone (**Fig. S6A-B**), we performed a similar analysis determine its relative contribution to location-specific MSN activity. In PRE sessions there was no difference in distribution of PYR cocaine index magnitudes (**Fig. S6C**), but in POST sessions the magnitude of the indices was larger for cocaine zone PYRs (*n* = 164, 205 PYRs, median 0.15 vs. 0.24; Wilcoxon rank sum test, *P* = 2.6 × 10^−4^, **Fig. S6D**), confirming that place cells over-represent the cocaine zone. Adjusting PYR spike trains to remove the asymmetry in PYR cocaine indices caused no change in MSN cocaine indices in PRE sessions (*n* = 531 MSNs, Wilcoxon signed rank test, *P* = 0.83, **Fig. S6E**), but in POST sessions the MSN indices showed a significant shift away from the cocaine zone (*n* = 713 MSNs, Wilcoxon signed rank test, *P* = 5.4 × 10^−4^, **Fig. S6F**), indicating that changes in PYR place field density also contribute to MSN recruitment to the cocaine zone. Since both place field changes and synaptic weight changes contributed significantly to increased MSN activity in the cocaine zone, we compared the changes directly to determine which was a larger contributor. In POST sessions, the two effects contributed equally to the location tuning properties of MSNs encoding the saline zone (*n* = 355 MSNs, Wilcoxon signed rank test, *P* = 0.29, **Fig. S6G**). However, for MSNs encoding the cocaine zone, changes in connections weights were a larger contributor than changes in PYR place fields (*n* = 340 MSNs, Wilcoxon signed rank test, *P* = 5.5 × 10^−9^, **Fig. S6H**).

## DISCUSSION

In this study, we found that cocaine place conditioning recruits NAc MSNs, and to a lesser extent hippocampal PYRs, to fire in the cocaine-paired zone. We also found that MSNs receive information about spatial location from PYRs, cocaine conditioning increases hippocampus-accumbens coupling, and MSNs modulated by the hippocampal theta rhythm show the strongest recruitment to the cocaine zone, all of which suggest that this effect is driven by hippocampal inputs to the NAc. Finally, we found that MSN recruitment is driven predominantly by preferential strengthening of hippocampal inputs encoding the cocaine zone.

Although extracellular recording is not able to make direct measurements of synaptic strength, ex vivo slice experiments have shown that repeated cocaine exposure potentiates hippocampal inputs to NAc MSNs (Britt et al., 2012; Pascoli et al., 2014). This provides the most likely cellular substrate for the selective increase in hippocampus-NAc coupling that we observe. Selective plasticity provides a possible explanation reconciling the conflicting results of Britt et al. (Britt et al., 2012), who found that cocaine selectively strengthens hippocampal inputs to the NAc, and MacAskill et al. (MacAskill et al., 2014), who found that cocaine selectively strengthens amygdalar inputs. Our results suggest instead that cocaine may preferentially strengthen the most active inputs, which could be predominantly hippocampal or amygdalar depending on the environmental conditions during cocaine exposure. It is worth emphasizing that our place conditioning paradigm (Fig. S2) pairs cocaine with a spatial location defined relative to distal navigational cues, while proximal sensory cues were minimized and counterbalanced across cocaine/saline conditions. Under similar conditions, CPP has been shown to be dependent on dorsal hippocampus and not ventral hippocampus or amygdala (Ferbinteanu and McDonald, 2001; Meyers et al., 2003; Trouche et al., 2016). In CPP paradigms incorporating proximal sensory cues that differ between the cocaine and saline zones, circuits beyond dorsal HPC are likely recruited as well.

Our results appear to contradict those of German et al. (German and Fields, 2007), who recorded neurons in the NAc during CPP POST sessions in rats and found that NAc MSNs exhibit decreased firing rates in the drug-paired zone. A possible explanation for this difference is that their study used morphine, and ours used cocaine. An interesting paradoxical observation is that both morphine and cocaine produce CPP, self-administration, and other addiction-related behaviors, but cocaine exposure increases the density of dendritic spines in the NAc and morphine decreases it (Robinson and Kolb, 1999a, b). This paradox was addressed recently by Graziane et al. (Graziane et al., 2016), who found that cocaine exposure forms new spines selectively on D1-positive MSNs, while morphine exposure prunes spines selectively on D2-positive MSNs. Given the previous observation that cocaine selectively potentiates hippocampal inputs onto D1-positive MSNs in the NAc (Pascoli et al., 2014), it is tempting to speculate that the increased firing we observed in the cocaine-paired zone is attributable to D1-positive MSNs, while the decreased firing German et al. observed in the morphine-paired zone is attributable to D2-positive MSNs. A recent study by Calipari et al. (Calipari et al., 2016) using fiber photometry provides some support for this hypothesis, but resolution of this issue will ultimately require future studies combining dual site unit recordings in the hippocampus and NAc with optogenetic tagging (Lima et al., 2009) of D1- or D2-positive MSNs.

Selective plasticity supports a plausible mechanistic model of cocaine CPP in which the NAc acts as an action-location-outcome associator (Berke and Hyman, 2000; van der Meer and Redish, 2011b). We hypothesize that the NAc integrates hippocampal inputs encoding a spatial location and prefrontal cortical inputs encoding an action plan to generate actions appropriate for a given spatial context. Under physiological conditions, performing an action in a specific location that generates a positive reward prediction error causes dopamine release in the NAc (Hart et al., 2014). This would strengthen hippocampal synapses encoding the rewarded location (Reynolds and Wickens, 2002), increasing the expected reward outcome associated with that action/location pairing. During cocaine place conditioning, NAc dopamine levels are artificially elevated, effectively rewarding all actions performed in the cocaine zone and increasing the strength of hippocampal inputs encoding that location. Upon exposure to the cocaine zone in POST sessions, the strengthened hippocampal inputs would drive increased firing in NAc MSNs, leading the animal to engage in activity in the cocaine zone rather than run to the saline zone. Determining whether this model is correct will require further studies involving multisite unit recording and manipulation of prefrontal cortex in CPP.

It is believed that drugs of abuse exert differential effects at various synaptic pathways in the brain – our results suggest that synaptic changes can be specific even at the level of functionally defined subsets of synapses within a single synaptic pathway. This reveals an additional layer of complexity that is not recognized in many circuit models of drug addiction and suggests that some of these models may require revision. Selective plasticity of specific corticostriatal synapses based on presynaptic firing properties has been observed in other striatal regions (Xiong et al., 2015) and may represent a canonical mechanism by which sensory and contextual information can bias action selection based on reward history.

## AUTHOR CONTRIBUTIONS

Conceptualization, L.S. and G.B. Investigation, L.S., A.C., and D.C. Formal Analysis, L.S., A.P., and A.C. Writing, L.S. and G.B.

## ACKNOWLEDGMENTS

The authors thank Gord Fishell, Lisa Roux, Viola Woo, and the members of the Buzsáki lab for support and comments on the manuscript. This work was supported by the Leon Levy Foundation (L.S.), the Brain and Behavior Research Foundation (L.S.), and NIH grants K08 DA036657 (L.S.) and MH107396 (G.B.). The authors have no conflicts of interest to report.

